# Antigen accumulation in the B cell follicle is impaired in aged mice

**DOI:** 10.64898/2026.01.30.702770

**Authors:** Xin Ge, Sigrid Fra-Bido, Silvia Innocentin, Hanneke Okkenhaug, Michelle A. Linterman, Louise M. C. Webb

**Author notes:** Co-senior authors (Michelle A. Linterman, Louise M. C. Webb).

## Abstract

Vaccine responses are impaired in aged mice and older people. An essential early step in mounting successful responses to vaccines is the delivery of antigens from the site of injection to the draining lymph node. This study tested the hypothesis that particle drainage to the lymph node is impaired in older mice after intramuscular injection. We observed fewer 20nm nanoparticles in the B cell follicle of the draining lymph node in aged mice after intramuscular injection. However, there was no significant difference in the abundance of larger 1000nm particles in the lymph nodes of aged mice, which are typically transported to the lymph node by antigen presenting cells. This data suggests that access of smaller antigens to the B cell follicle is impaired in ageing and may have implications for the initiation of T and B cell responses after vaccination.

## Introduction

Vaccination provides the body with a small dose of non-pathogenic antigen that stimulates the immune system to generate antibodies and memory T and B cells that are specific for the invading pathogens, facilitating protection against future encounters. However, ageing of the mammalian immune system contributes to a decline in vaccine efficacy. Vaccination induces the humoral and the cell-mediated immune responses, with humoral immunity being negatively impacted by ageing across multiple vaccine types (Lee & Linterman, 2022). Humoral immunity is generated via two pathways: First, the extrafollicular responses that takes place rapidly after immunisation (Eisenbarth et al., 2025), and second, via the germinal centre response that gives rises to high-affinity antibodies where B cells undergo clonal expansion, mutation and selection to produce long-lived memory B cells and antibody secreting cells (De Silva & Klein, 2015). Whilst the extrafollicular response is intact in aged mice (Silva-Cayetano et al., 2021), the initiation, size and quality of the germinal centre responses is significantly impaired with advancing age (Luscieti et al., 1980; Silva-Cayetano et al., 2023; Szakal et al., 1990). B cell extrinsic factors contribute to the poor germinal centre responses seen in aged mice (Denton et al., 2022; Foster et al., 2025; Lee et al., 2022; Lee et al., 2023). However, the full extent of these age-dependent changes has not been established.

Lymph nodes are key hubs for initiating humoral immunity after vaccination, bringing together rare antigen-specific lymphocytes with antigens from peripheral tissues. Following vaccination, antigens travel from the local site of injection to the lymph node through lymphatic vessels, either via free drainage in the lymphatic fluid, or by transportation of antigen presenting cells that have acquired the antigen within tissues (Randolph et al., 2017). Antigens delivered to the lymph node can then trigger immune responses when they encounter and subsequently activate cognate T and B cells in the presence of co-stimulatory factors.

Several studies have suggested that antigen transport in the lymph may be defective in the context of advanced age. Rodent (Ahn et al., 2019; Akl et al., 2011; Antila et al., 2017; Da Mesquita et al., 2018; Gasheva et al., 2007; Karaman et al., 2015; Kataru et al., 2022; Lei et al., 2025; Rustenhoven et al., 2023; Zolla et al., 2015), non-human primate (Thompson et al., 2019) and human (Albayram et al., 2022; Conway et al., 2009; Donato et al., 2009; Unno et al., 2011; Zhou et al., 2020) studies consistently report declining lymphatic function in the later years of life. Before reaching the lymph node, antigens from the interstitial fluid travel through lymphatic capillaries and lymphatic vessels. The contraction of the muscles surrounding the lymphatic vessels support the flow of the lymph from local tissue towards the lymph node (Randolph et al., 2017). Contractility of lymphatic vessels impacts lymphatic drainage (Zolla et al., 2015). In addition, with age, lymphatic networks display reduced density, impaired contractility and increased leakiness (Ahn et al., 2019; Karaman et al., 2015; Zolla et al., 2015) As the lymph brings various antigens from the peripheral tissues into the lymph node, impaired lymphatic function may cause limited antigen availability in the lymph node. In the meningeal lymphatics that drain the central nervous system, a defect of antigen drainage to the brain is reported in aged individuals causing neuroinflammation and reduced cognition (Ahn et al., 2019; Bolte et al., 2020). This suggests that there are age-associated changes in lymphatic biology that affect antigen access to the lymph node after vaccination.

In this study, we set out to test the hypothesis that antigen drainage to the lymph node is impaired in ageing after intramuscular injection. To test this, we imaged the draining lymph nodes of younger adult (9-19 weeks-old) and aged (95-100 weeks-old) mice following injection of fluorescent polystyrene nanoparticles of specific sizes to explore age-associated changes of the location and quantity of antigens in the lymph node upon vaccination. Two sizes of fluorescent nanoparticles (20nm and 1000nm) were used to distinguish free drainage (20nm) from dendritic cell-mediated antigen transport (1000nm) (Manolova et al., 2008). Two days after intramuscular injection, we observed fewer 20nm particles in the B cell follicle of lymph node in aged mice, but no significant difference in the abundance of 1000nm particles in the lymph nodes. This indicates that the availability of smaller particles in the lymph node may be impaired after intramuscular injection.

## Results

### Fluorescent nanoparticles are present in the medial iliac lymph node of both adult (9-19 weeks) and aged (95-100 weeks) mice after intramuscular immunisation

Adult (9-19 weeks) and aged (95-100 weeks) mice were injected intramuscularly in the right quadricep with 20nm and 1000nm fluorescent polystyrene nanoparticles. Two days after immunisation, draining medial iliac lymph nodes were imaged to determine the location and abundance of the nanoparticles (Fig 1A). The size of the nanoparticles has been previously shown to determine the mode of transport into the lymph node with 20nm particles reported to drain freely in the lymph to the lymph node, and 1000nm red particles transported by antigen presenting cells (Manolova et al., 2008). Two days after nanoparticle injection, the medial iliac lymph nodes were taken from both adult (9-19 weeks) and aged (95-100 weeks) mice and co-stained for IgD (Fig 1B) and CD3 (Fig 1C) separately, to facilitate identification of the T and B cell areas within the lymph node. The CD3+ T cell zone is in the lymph node paracortex and surrounded by IgD+ B cell follicles which are beneath the subcapsular sinus at the outer cortical layer of the lymph node. The staining of IgD and CD3 is consistent with the typical lymph node structure in both 9-19 weeks old and aged 95-100 weeks old mice (Girard et al., 2012).

**Figure 1.**
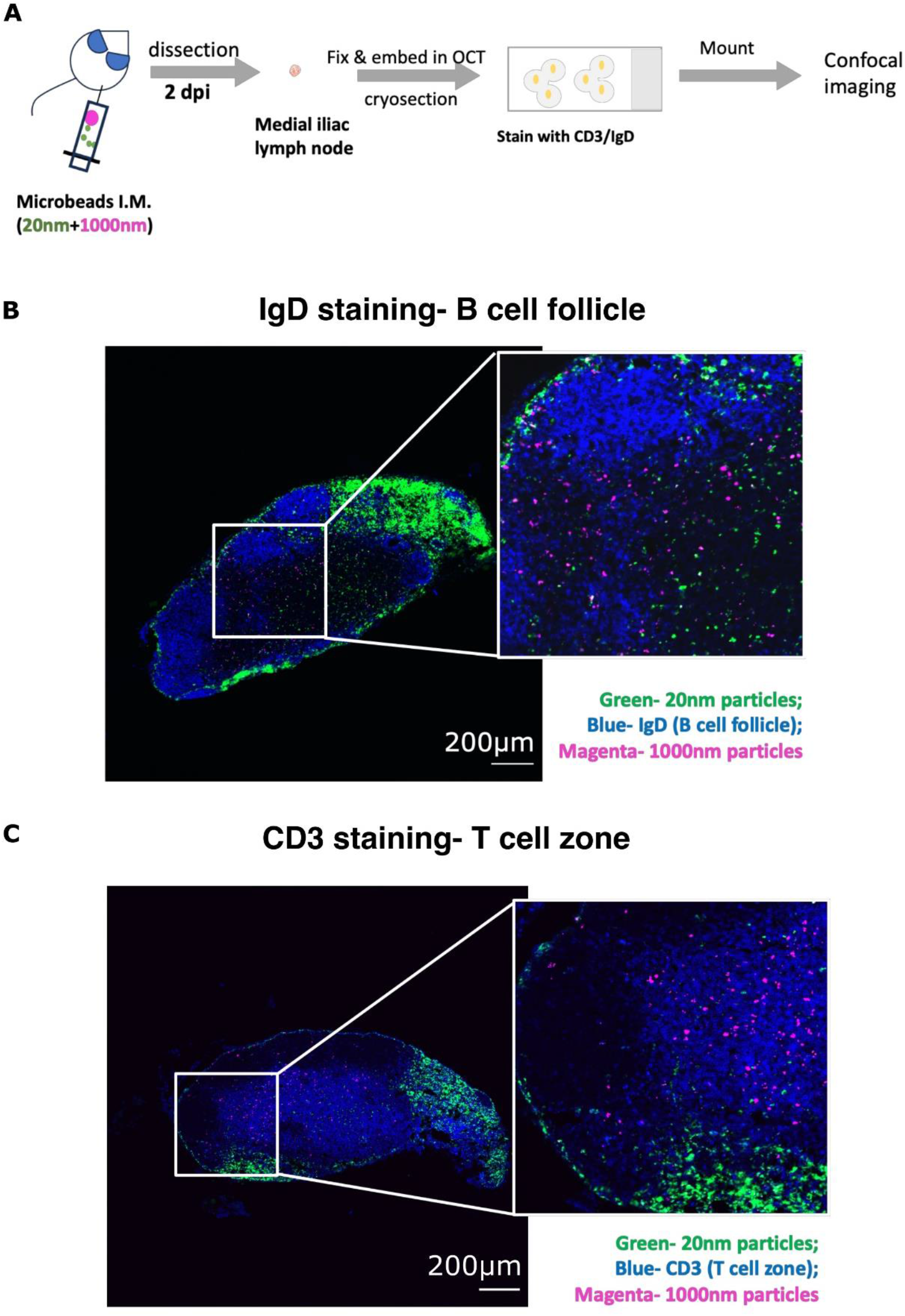
Medial iliac lymph nodes two days post nanoparticle injection. A) Experimental design and workflow for intramuscular immunisation of 20nm and 1000nm particles, draining lymph node collection and imaging. B) Example staining of the B cell follicle (IgD) in medial iliac lymph nodes (draining lymph node). C) Example staining of the T cell zone (CD3) in medial iliac lymph nodes (draining lymph node).

Confocal imaging confirmed the presence of both 20nm and 1000nm nanoparticles in the lymph node two days after injection. Both sizes of particles formed small clusters in the lymph node, since the observed signal is irregularly shaped and mostly larger in size than the diameter of the nanoparticles (Fig 1B and C). Two days after co-injection, most of the red 1000nm nanoparticles appeared to be in the T cell zone, while the majority of green 20nm nanoparticles accumulated at the outer border of the B cell follicle.

### Image quantification reveals regional richness of nanoparticles in adult (9-19 weeks) and aged (95-100 weeks) draining lymph nodes

To determine whether age altered the location and number of fluorescent nanoparticles we developed an automated quantification pipeline with CellProfiler™ (Jones et al., 2008) (Fig 2A). The pipeline identifies and segments CD3^+^ and IgD^+^ areas, as well as and the whole lymph node (Fig 2B and C). After area segmentation, the fluorescent signal within the collected spectrum range (638-756nm for 1000nm particles and 499-624 nm for 20nm particles) of the 20nm (Fig 2D) and 1000nm (Fig 2E) particles were identified separately based on geometric pattern recognition that distinguishes nanoparticles from irregularly shaped background noise, and then intensity level thresholding that further distinguishes nanoparticles from the low-intensity background.

**Figure 2.**
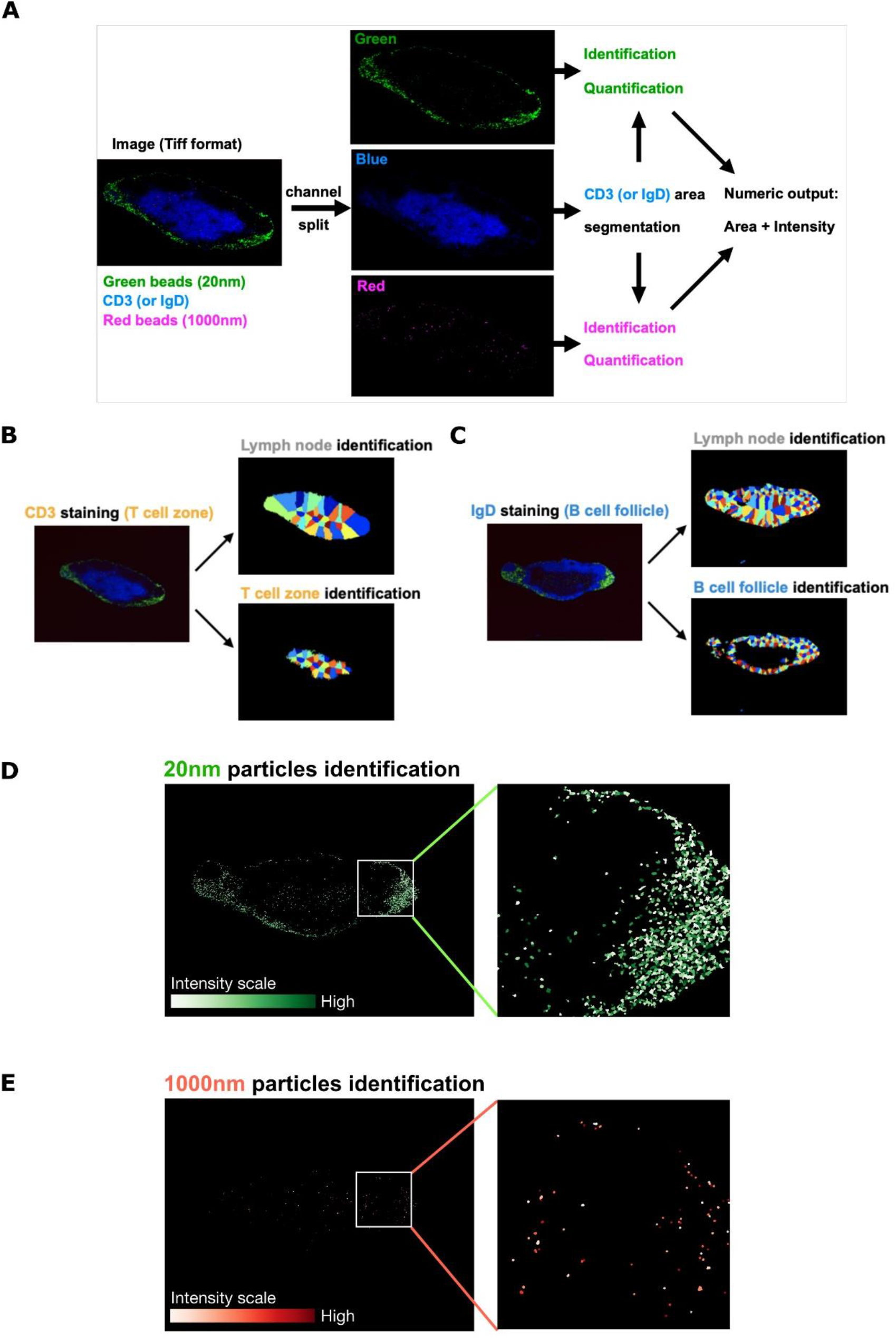
Pipeline workflow designed to reveal identification and quantification of areas of the lymph node and the nanoparticles. A) Quantification pipeline designed to measure the abundancy of both 20nm and 1000nm particles in CD3+ T cell zone, IgD+ B cell follicle node, or the whole lymph node. B-C) Automatic segmentation of CD3+ T cell zone (B) or IgD+ B cell follicle (C) and whole lymph node. D) Identification of 20nm particles in segmented regions, original and zoomed view. White-green colormap for intensity scale. E) Identification of 1000nm particles in segmented regions, original and zoomed view. White-red colormap for intensity scale.

### Fewer 20nm nanoparticles are observed in lymph nodes from aged (95-100 weeks) mice compared with adult (9-19 weeks) animals

20nm and 1000nm nanoparticles were quantified in two ways: the occupied area in pixels and total intensity value of the identified pixels. This was done in the segmented regions including T cell zone, B cell follicle and the whole lymph node, both with and without normalisation to the total area of the specified region (Table 1). With image input into the pipeline, followed by statistical analysis of pipeline output values, we could quantify the total area of the IgD^+^ and CD3^+^ regions and the abundance of nanoparticles within these areas.

**Table 1.** Output parameters of quantification pipeline. Measurements of area of segmented region as well as area occupied and total intensity of microbeads in segmented region calculated by the pipeline. LN = lymph node, CD3 area = CD3 labelled T cell zone, IgD area = IgD labelled B cell follicle.

| Output parameters | Explanation |
| --- | --- |
| CD3 area (or IgD area) | Area by pixel of identified CD3/IgD area |
| LN area | Area by pixel of identified LN area |
| Area green in CD3 (or IgD) | Area by pixel of identified green beads in CD3/IgD area |
| Area green in LN | Area by pixel of identified green beads in LN area |
| Area red in CD3 (or IgD) | Area by pixel of identified red beads in CD3/IgD area |
| Area red in LN | Area by pixel of identified red beads in LN area |
| Intensity green in CD3 (or IgD) | Total intensity of identified green beads in CD3/IgD area |
| Intensity green in LN | Total intensity of identified green beads in LN area |
| Intensity red in CD3 (or IgD) | Total intensity of identified red beads in CD3/IgD area |
| Intensity red in LN | Total intensity of identified red beads in LN area |

The quantification pipeline (Fig 2, Table 1) was first applied to images of lymph nodes taken two days after nanoparticle injection that are co-stained with IgD. The total area of the lymph node section was not different between age groups (Fig 3A, B), but the 20nm nanoparticles were less abundant in the aged lymph node compared with their younger adult controls (Fig 3C). By contrast, the abundance of the 1000nm nanoparticles was not different between age groups. (Fig 3D). The area of the IgD^+^ B cell follicle was comparable between age groups (Fig 3E). In the B cell follicle area, the normalised intensity of the 20nm nanoparticles was significantly lower in aged mice compared to their younger adult controls (Fig 3F). But the amount of the 1000nm particle in the B cell follicle did not change with age, consistent with what has just been observed in the whole lymph node (Fig 3G, Fig 3D). This indicated that there is a size-dependent impairment in antigen access to the B cell follicle in 95-100-week-old animals after nanoparticle injection.

**Figure 3.**
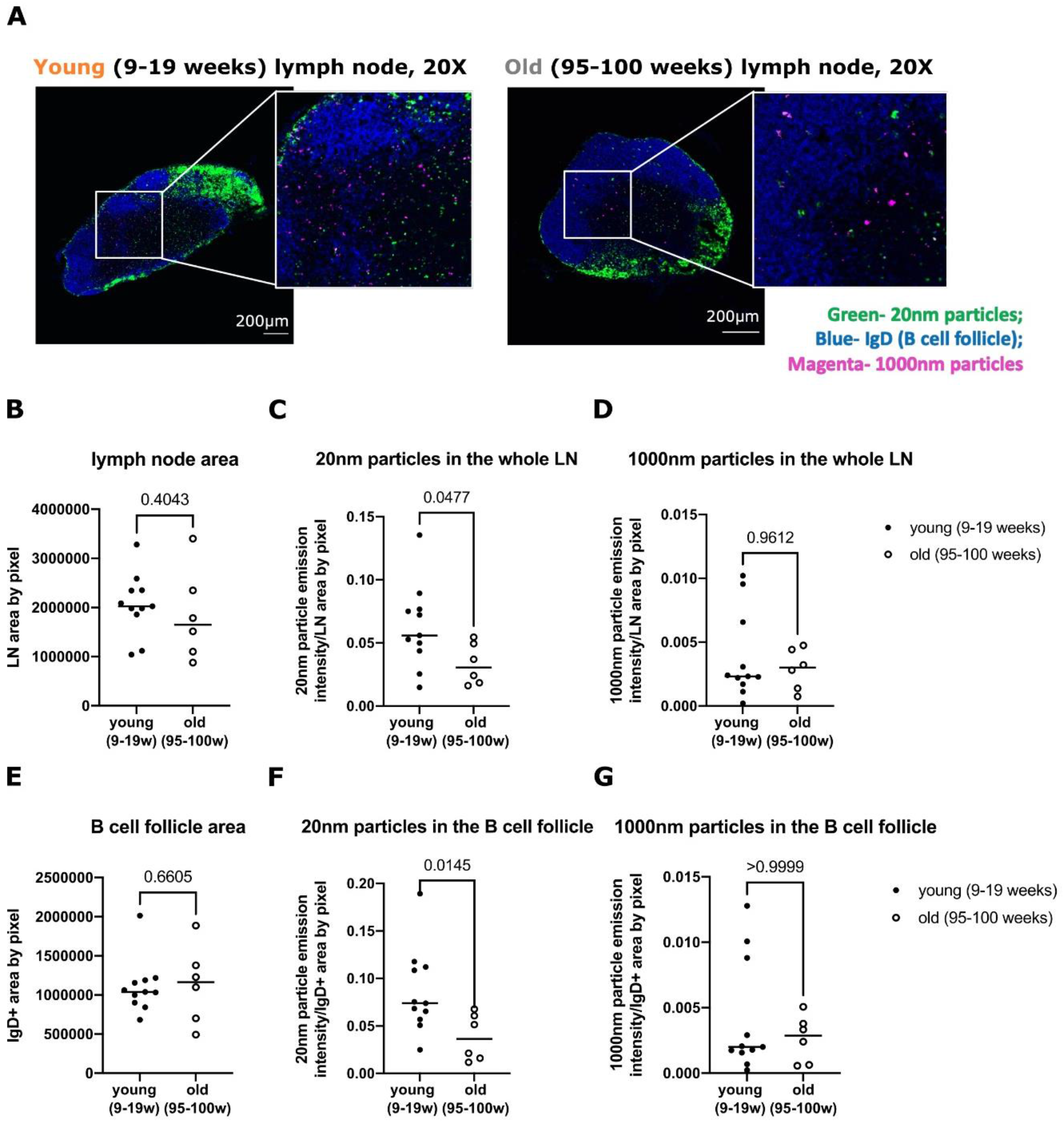
Quantification of the whole draining lymph node and the B cell follicle 2 days post intramuscular immunisation in aged mice. A) Example images of young (9-19 weeks) and aged (95-100 weeks) lymph node section with IgD staining for the B cell follicle. B) Quantified area by pixel of lymph node section in young (9-19 weeks) vs aged (95-100 weeks) mice. C) Normalised intensity of 20nm particles in the whole lymph node. D) Normalised intensity of 1000nm particles in the whole lymph node. E) Quantified area of the B cell follicle by pixel. F) Normalised intensity of 20nm particles in the B cell follicle. G) Normalised intensity of 1000nm particles in the B cell follicle. Each symbol represents one biological replicate, median shown for each group. Mann Whitney test carried out to examine difference between groups. Experiment repeated twice with one representative experiment shown.

### Comparable abundance of 20nm and 1000nm nanoparticles in the T cell zone across age groups

To understand whether the effect observed in the whole lymph node is purely from the B cell follicle, or if nanoparticle access to the T cell zone is also altered in ageing, we applied the quantification pipeline to the CD3^+^ sections of the same draining lymph nodes two days post injection of 20nm and 1000nm nanoparticles. The area of the CD3^+^ T cell zone was reduced in the aged compared to younger adult mice (Fig 4A, B). This has been observed in a replicate experiment (*p* = 0.0549; Supplementary Table 1). The normalised intensity of the 20nm particles (Fig 4C) and 1000nm particles in the T cell area (Fig 4D), were not found to be significantly different between young and aged mice in both experiments (Fig 4C, D, Supplementary Table 1). Altogether, the above data suggests that access of the 20nm nanoparticle to the B cell follicle is diminished in aged mice, while no age-dependent changes in the abundance of 1000nm nanoparticle access was detected.

**Figure 4.**
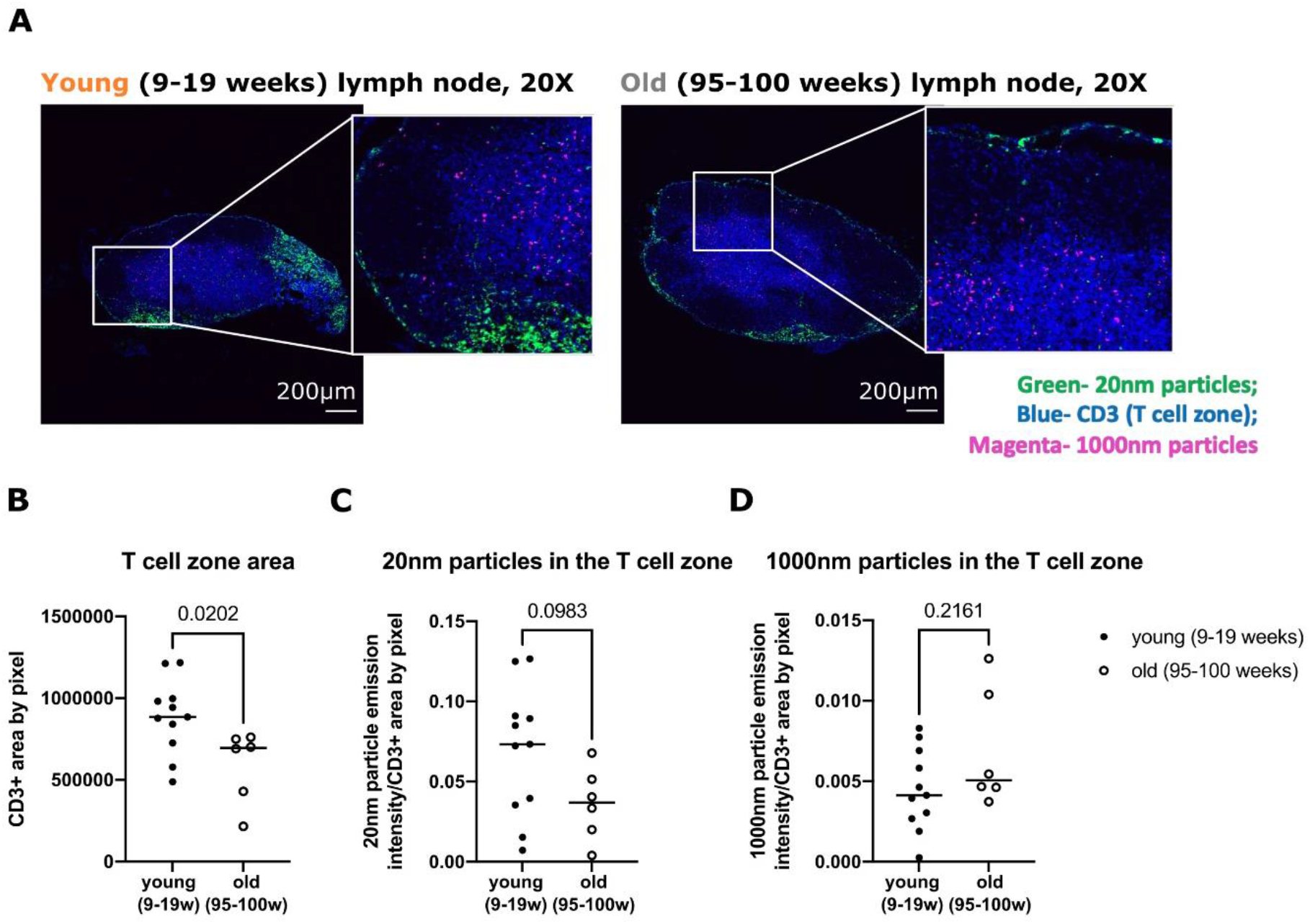
Quantification of the T cell zone 2 days post intramuscular immunisation in aged mice. A) Example image of young (9-19 weeks) and aged (95-100 weeks) lymph node section with CD3 staining for the T cell zone. B) Quantified area by pixel of the T cell zone in young (9-19 weeks) vs aged (95-100 weeks) mice. C) Normalised intensity of 20nm particles in the T cell zone. D) Normalised intensity of 1000nm particles in the T cell zone. Each symbol represents one biological replicate, median shown for each group. Mann Whitney test carried out to examine difference between groups. Experiment repeated twice with one representative experiment shown.

## Discussion

In this study, we utilised fluorescent nanoparticles of two different sizes to assess antigen access to the lymph node after intramuscular injection. Free antigens access the lymph node via the lymph flow, and consistent with this we observed the 20nm particles at the edge of the lymph node and in the B cell follicle. The 20nm nanoparticles were less abundant in the B cell follicle of 95-100 weeks aged mice compared with 9-19 weeks adult mice. By contrast, the abundance of the 1000nm particles, which are thought to be transported by antigen presenting cells into the lymph node, was not altered in ageing. Together this suggests that drainage of particles in the lymph fluid to the lymph node may be impeded by older age.

Antigens travel from local tissue to the lymph node via the lymphatics before being able to trigger downstream immune responses (Randolph et al., 2017). It has been previously reported that the rate of lymph flow is reduced in aged rats (Zolla et al., 2015). It is also known that the immune responses in secondary lymphoid organs decline with age (Foster et al., 2025). To our knowledge, no direct relationship has been established between ageing and antigen availability in the lymph node after intramuscular injection. Our finding that fewer 20nm particles are observed in B cell follicles of aged lymph nodes indicates that ageing negatively affects the availability of antigens in the lymph node.

The method of transport for lymph-carried antigens is size-dependent. In our immunisation model, we injected nanoparticles of two different sizes, 20nm and 1000nm, to assess antigen free drainage and dendritic cell mediated carriage, respectively (Manolova et al., 2008). Upon reaching the lymph node, smaller sized antigens (<70 kDa; for example, 20nm nanoparticles in our experiment) drain passively into the subcapsular sinus, after which they enter lymph node parenchyma via reticular conduits and can then be captured by local B cells, dendritic cells and macrophages (Itano et al., 2003). Antigen presentation by macrophages have been reported to decline with age (Herrero et al., 2001). However, the 20nm polystyrene particles used in our study, are unlikely to be immunogenic, suggesting that the age-associated decrease in 20nm particles observed in the B cell follicle reflects defective passive antigen drainage rather than downstream antigen processing by antigen-presenting cells in the lymph node of older mice.

The 1000nm particles, unable to drain freely into the lymph node due to their large size, are reported to rely on dendritic cell mediated carriage to enter the lymph node. Conventional dendritic cells type II (cDC2) have been implicated in T follicular helper cell priming as the dominant DC subset early after immunisation (Krishnaswamy et al., 2018). It has been observed that aged mice have fewer antigen-bearing cDC2 in the draining lymph node after immunisation, and these cells have reduced expression of peptide-MHC II and costimulatory receptors than cDC2 from younger adult mice (Stebegg et al., 2020). However, in our non-immunogenic particle model, no difference of DC-dependent carriage was observed. This is likely because the nanoparticles used are not immunogenic and suggests that the age-associated deficiencies seen in other studies reflects impaired dendritic cell activation signalling triggered by immunogenic antigens.

The efficacy of various commonly used vaccines has been found to decline with age. Seasonal influenza vaccines, for example, elicit a lower titre antibody response in the elderly (age >60 years of age) compared with younger adults (age <65 or <60, varies for different studies) as described in a systemic review of 31 vaccine antibody response studies conducted from year 1986 to 2002 (Goodwin et al., 2006). The primary antigen response to yellow fever vaccination has been reported to alter in ageing, reflecting in a delay in antibody production (Roukens et al., 2011). This delay in antibody response could be explained by limited antigen availability in the B cell follicle of the elderly as observed in our study. As the site of B cell activation, reduced antigen availability in B cell follicles in the elderly may contribute to a declined B cell activation and thus a declined scale of the germinal centre response and antibody production (Silva-Cayetano et al., 2023).

Antigen availability remains a key factor in vaccine design and several approaches has been developed to improve vaccine responses in the elderly. One of the strategies is to give aged individuals repeated dose of vaccination. A recent study has shown that compared with a single dose, two doses of ChAdOx1 nCoV-19, a vector-based COVID vaccine, gives more protection in older people (Paranthaman et al., 2022). Specific adjuvants in vaccination are also developed to stimulate the aged immune system (Nanishi et al., 2022). For example, oil-in-water emulsion adjuvant MF59 and AS03, as well as liposome-based adjuvant AS01 has been applied in influenza vaccines to raise vaccine efficacy in elder population (Wagner & Weinberger, 2020). Another strategy that is consistent with the findings in our study is to use a higher dose of vaccine antigens. For example, high-dose influenza vaccine has been used to improve protection and reduce mortality among older adults (Chaves et al., 2023). The 20nm nanoparticles used in this study are of similar size to viral particles and protein subunits typically used for vaccination (Pollard & Bijker, 2021). The reduced availability of the 20nm particles in the B cell follicle of aged lymph nodes suggests that increased dosing of viral particle or protein/polysaccharide subunit vaccines could be considered for vaccination of older persons to circumvent the defects in antigen drainage to the lymph node enabling immune response elicitation.

## Materials and Methods

### Mice

Wild type (WT) C57BL/6 mice (strain C57BL/6Babr) were bred, housed, and aged in the Babraham Institute Biological Support Unit under specific pathogen-free conditions. Housing with ambient temperature of 19–21°C and relative humidity 52% was provided. Lighting of 12 hours of day and 12 hours of night was monitored, with 15mins of transition period with moderate lighting inclusive. All cages were individually ventilated and received environmental enrichment including nesting material, plastic houses and chew sticks, holding 1-5 per cage for mice after weaning. Nourishment supply includes CRM (P) VP diet (Special Diet Services) ad libitum as main diet plus seeds (eg. sunflower, millet) as cage-cleaning-period enrichment. All mouse experimentation was performed with the approval of the Babraham Institute Animal Welfare and Ethical Review Body. Animal husbandry and experimentation were all done in accordance with existing European Union and United Kingdom Home Office legislation and local standards under the home office licence P4D4AF812. Young WT C57BL/6 mice were 8-19 weeks old and aged WT C57BL/6 mice were 95-100 weeks old when experiments were started. All experiments were performed according to the regulations of the UK Home Office Scientific Procedures Act (1986) under the PPL P4D4AF812.

### Immunisation

For administration of nanoparticles, mice were injected intramuscularly in the right quadricep with 50μl of 1:1:4 mixture of yellow-green fluorescent (505/515) FluoSpheresTM Carboxylate-modified Microspheres (ThermoFisher Scientific #F8787, referred to as 20nm green nanoparticles), crimson fluorescent (625/645) FluoSpheresTM Carboxylate-modified Microspheres (ThermoFisher Scientific #F8816, referred to as 1000nm red nanoparticles) and Phosphate-Buffered Saline (PBS).

### Dissection

Mice were euthanized and medial iliac lymph nodes, located at the branch point of the aorta and the iliac vessel, were dissected and placed into PBS for preparation for microscopy.

### Immunofluorescence and confocal microscopy

Lymph nodes were immersion-fixed in diluted BD Cytofix (BD Biosciences #554722, diluted 1:3 with PBS) at 4°C for 1-3 days, washed in PBS and incubated in 30% sucrose at 4°C for 1-3 days for cryoprotection. Fixed and cryoprotected tissue were embedded in optimal cutting temperature (O.C.T.) medium (Scigen #23-730-625 or Fisher HealthCare #23-730-571) in 2-propanol bath on dry ice and stored at -80°C. Frozen tissues were cut into 10μm sections at -20°C on cryostat (Leica CM1850). Sections were air-dried at room temperature and stored at -20°C (short term) or -80°C (long term). For staining, sections were defrosted and air-dried at room temperature (RT), enclosed in hydrophobic circle drawn by ImmEdge Pen (Vector laboratories #H-4000), rehydrated with wash buffer, blocked in wash buffer containing 2% BSA (Jackson ImmunoResearch #001-000-161) for 1hr at RT and permeabilised with 1% Triton X-100 (Sigma #X100, diluted in PBS) for 30mins at RT. Tissue sections were stained either with eFluor 450 conjugated anti-IgD antibody (11-26c, eBioscience #48-5993-82, 1:200 dilution with wash buffer containing 1% BSA) or Pacific Blue conjugated anti-CD3 antibody (17A2, BioLegend #100214, 1:200 dilution with wash buffer containing 1% BSA) overnight at 4°C. Sections were washed once with wash buffer and then twice with PBS, mounted in aqueous mounting media (National Diagnostics Hydromount #HS-106), covered gently with coverslips (22x22mm, 0.16-0.19mm thick) avoiding trapping of air bubbles, and dried at RT for at least 30mins. Mounted sections were imaged with Zeiss LSM780 confocal microscope and Zen black edition software (Zeiss) using 405nm (Diode 405-30), 488nm (Argon) and 633nm (HeNe633) lasers and 20x/0.8 air objective (Zeiss Plan-Apocheromat 20x/0.8 M27), pixel size of 0.69μm, and a zoom of 0.6. Tissue sections were imaged using the tile scan mode to acquire large field of view of the whole lymph node, 5% overlap of tiles. Blue (405nm laser, Ch1 detector, signal range 424-502nm) and red (633nm laser, Ch2 detector, signal range 638-756nm) channels are taken in the same track while separate from green channels (488nm laser, ChS1 detector, signal range 499-624 nm). At least 6 sections for both CD3 and IgD staining were taken and imaged per mouse.

### Image quantification

Area of the T cell zone, B cell follicle and the whole lymph node section by pixel as well as the area and intensity of two types of microbeads in specified regions (either with or without normalisation by area of specified region) were calculated in Cell Profiler with an automated pipeline developed by Sigrid Frá-Bido for identification of specified region as well as identification and quantification of the microbeads. ∼4 sections on average per mouse were qualified for quantification and analysed via the pipeline. Images of the original CZI format were converted to TIFF format before being loaded to the pipeline. Channels were split at the first step of analysis. CD3 and IgD staining in the blue channel were used to identify the T cell zone and B cell follicle respectively. After identification of primary objects (T cell zone and B cell follicles), the whole lymph node region was identified based on the primary objects. Microbeads were then identified during which signal in green and red channels were enhanced based on geometric pattern and automatic thresholding (global thresholding with Minimum Cross-Entropy method) was set up based on intensity. The area and intensity of identified objects (20nm and 1000nm particles) in specified regions (T cell zone or B cell follicle and the whole lymph node) were calculated, and the values were summarised in an output CSV file. Bulk analysis of images with the Cell Profiler pipeline was carried out via high performance computing with the Babraham computing cluster. Normalisation was done through dividing the area (in pixel) and intensity value of microbeads in specified regions by the area (in pixel) of the regions afterwards.

Values from image quantification were combined by sample via averaging output value of multiple images from the same mice. Prism 9 (GraphPad) software was use for analysis and visualisation of combined data. Statistical significance was determined using Mann-Whitney test where young and aged groups are compared on the same time point. P-values <0.05 were considered as statistically significant.

## Author Contributions and Notes

Xin Ge carried out mouse experiments, image collection and image analysis on local devices and the cluster. Sigrid Fra-Bido designed and optimised the image analysis pipeline. Silvia Innocentin helped with mouse experiments, mouse colony management, preparation of beads for injection and tissue collection. Hanneke Okkenhaug assisted with finalising the image analysis pipeline and provided Xin Ge training on cluster usage for image analysis. Michelle Linterman and Louise Webb provided guidance and supervision on experiment design and data analysis.

## Acknowledgments

We acknowledge the contribution of the Babraham Institute Biological Support Unit staff, who performed in vivo treatments of our animals and took care of animal husbandry. We thank the staff of the Babraham Imaging facility for their technical support. This study was supported by funding from the Lister Institute of Preventative Medicine, Biotechnology and Biological Sciences Research Council (BBS/E/B/000C0427 to M.A.L. and the Campus Capability Core Grant to the Babraham Institute). This research was conducted while X.G. was a PhD student and the author gratefully acknowledge the PhD studentship supported by the Cambridge Commonwealth, European and International Trust and the China Scholarship Council.

