## Supplementary material for "Antigen accumulation in the B cell follicle is impaired in aged mice": Supplemenrary Table and Figures

**Supplementary Table 1. Statistics for repeat experiment.** Repeat experiment with young (12-13 weeks, n=9) vs old (96-97 weeks, n=7) mice day 2 post immunisation. Statistics done with Mann-Whitney test.

| Parameter | Trend | p value |
|---|---|---|
| LN area | Young > old | 0.1738 |
| IgD area | None | 0.7577 |
| CD3 area | Young > old | 0.0549 |
| Norm intensity 20nm in LN | Young > old | 0.0907 |
| Norm intensity 20nm in IgD | Young > old | 0.0549 |
| Norm intensity 20nm in CD3 | None | 0.4698 |
| Norm intensity 1000nm in LN | None | 0.8371 |
| Norm intensity 1000nm in IgD | None | 0.9182 |
| Norm intensity 1000nm in CD3 | None | >0.9999 |

**A**

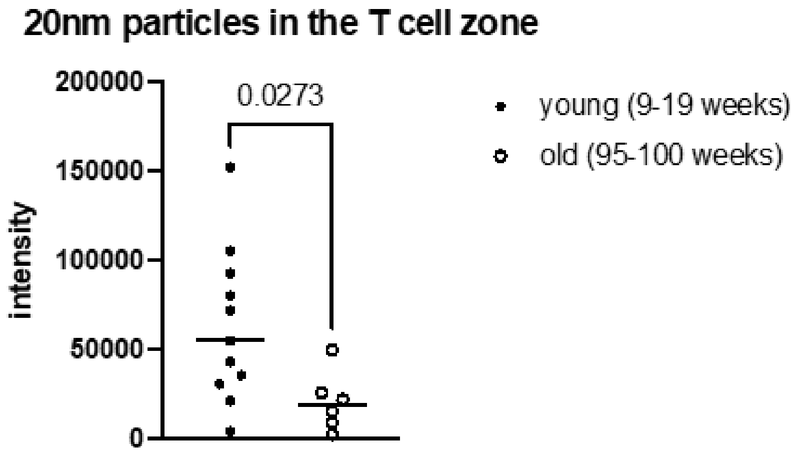


**B**

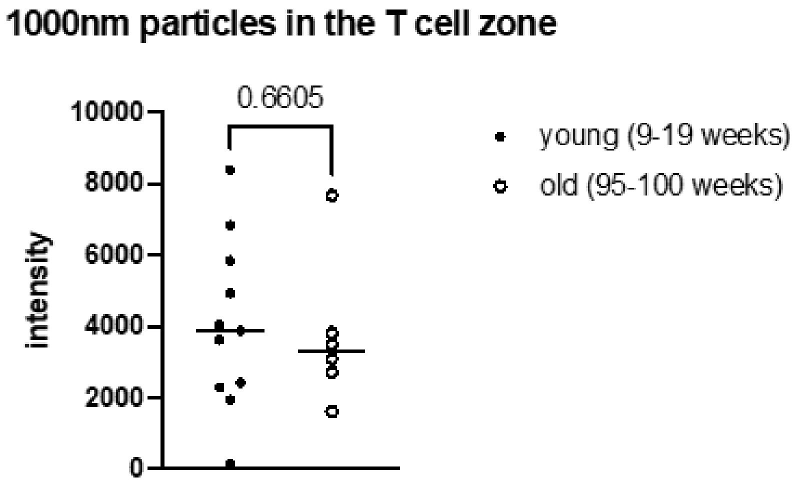


**Supplementary Figure 1. Unnormalised intensity of 20nm and 1000nm particles in the T cell zone.** A) Unnormalised intensity of 20nm particles in the T cell zone. B) Unnormalised intensity of 1000nm particles in the T cell zone. Each symbol represents one biological replicate, median shown for each group. Mann Whitney test carried out to examine difference between groups. Experiment repeated twice with one representative experiment shown.
